# Calibration of consonant perception to room reverberation

**DOI:** 10.1101/2020.09.01.277590

**Authors:** Eleni Vlahou, Kanako Ueno, Barbara G. Shinn-Cunningham, Norbert Kopčo

## Abstract

**Purpose:** We examined how consonant perception is affected by a preceding speech carrier simulated in the same or a different room, for a broad range of consonants. Carrier room, carrier length, and carrier length/target room uncertainty were manipulated. A phonetic feature analysis tested which phonetic categories are most influenced by the acoustic context of the carrier.

**Method:** Two experiments were performed, each with 9 participants. Targets consisted of vowel-consonant (VC) syllables presented in one of 2 strongly reverberant rooms, preceded by a VC carrier presented either in the same room, a different reverberant room, or an anechoic room. In Experiment 1 the carrier length and the target room randomly varied from trial to trial while in Experiment 2 they were fixed within blocks of trials.

**Results:** Compared to the no-carrier condition, a consistent carrier provided only a small advantage for consonant perception, whereas inconsistent carriers disrupted performance significantly. For a different-room carrier, carrier length had an effect; performance dropped significantly in the 2-VC compared to the 4-VC carrier length. The only effect of carrier uncertainty was an overall drop in performance. Phonetic analysis showed that an inconsistent carrier significantly degraded identification of the manner of articulation, especially for stop consonants, and, in one of the rooms, also of voicing.

**Conclusions:** Calibration of consonant perception to strong reverberation is exhibited through disruptions in perception when the room is switched. The strength of calibration varies across different consonants and phonetic features, as well as across rooms and durations of exposure to a given room.

## Introduction

Reverberation is ubiquitous in everyday settings. It has a pervasive influence on the acoustic signal reaching a listener, affecting its temporal structure, spectral content, and interaural differences (Shinn-Cunningham, 2003). Numerous studies show that reverberation can impair spatial hearing and speech perception. For example, it negatively affects sound localization in the horizontal plane (Hartmann, 1983), selective auditory attention to a speech source in the presence of competing sources (Ruggles & Shinn-Cunningham, 2011), and speech intelligibility, particularly for children and older adults, nonnative listeners and hearing-impaired individuals (Assman & Summerfield, 2004; Lecumberri et al., 2010; Nábělek & Donahue, 1984; Takata & Nábělek, 1990). On the other hand, there is strong evidence that adult listeners can quickly adapt to and take advantage of reverberation in many situations (Helfer, 1994; Shinn-Cunningham, 2003). For instance, listeners are sensitive to the statistical regularities that are present in everyday reverberation and exploit these regularities to separate the contributions of sound sources and environmental filters (Traer & McDermott, 2016). Reverberation can facilitate distance perception (e.g., Zahorik, Brungart, & Bronkhorst, 2005). Furthermore, exposure to different rooms during phonetic training can enhance implicit phonetic learning (Vlahou et al., 2019). Collectively, these results suggest that reverberation can both disrupt and enhance auditory perception, and that listeners use various adaptation mechanisms to mitigate the negative impacts of reverberation and to improve auditory and speech perception.

Different researchers have postulated both monaural and binaural adaptation mechanisms that use information from preceding context to modify and improve speech perception in reverberation (Beeston et al., 2014; Brandewie & Zahorik, 2010; Srinivasan & Zahorik, 2013; Watkins, 2005). In a seminal study, Watkins (2005) exposed listeners to different levels of reverberation, using monaural speech tokens from the continuum from “sir” to “stir”. He showed that, for the same amount of reverberation imposed on the same speech token, listeners shifted their responses towards “sir” or “stir” depending on the level of reverberation in the preceding carrier phrase. Later studies replicated this finding with other speech sounds (Beeston et al., 2014) and non-speech contexts (Watkins & Makin, 2007). Zahorik and colleagues used binaural tasks with speech stimuli presented in reverberation and noise. They demonstrated that prior exposure to a consistent room significantly improved performance for stimuli taken from the Coordinate Response Measure corpus (Bolia et al., 2000; used in Brandewie & Zahorik, 2010) and for sentences with rich phonetic and lexical content taken from the TIMIT database (Garofolo et al., 1993; used in Srinivasan and Zahorik, 2013). These results provide robust evidence that exposure to consistent rooms improves subsequent speech processing, but also raise important new questions about what drives this perceptual recalibration.

First, while there is strong evidence that speech perception can be dramatically improved after exposure to consistent reverberation, less is known about how different inconsistent environments affect performance. Brandewie and Zahorik (2018, Experiment 1) replicated the finding of improved speech-in-noise perception after exposure to a consistent room, compared to a baseline condition where no prior room context was given. Examining the effects of inconsistent carriers, they found that when there was a switch from one reverberant room to a room with different reverberation, performance was significantly worse than in the consistent condition, and that the amount of degradation depended on the relative strength of reverberation in the carrier vs. target rooms. Specifically, the disruption was larger when the switch was from a more reverberant carrier room to a less reverberant target room, while it was smaller when the carrier room was less reverberant than the target room. The authors suggested that this might occur because some of the adaptation to the less-reverberant carrier transferred to the new room, improving performance and attenuating the difference with the consistent condition. Importantly, the disruption never resulted in performance worse than the no-carrier baseline. These results motivate further examination of how the acoustic properties of a preceding and new environment interact, especially when the speech is not masked by noise and the system can tune itself optimally to the room characteristics. Specifically, we hypothesize that when examined in strongly reverberant rooms without noise masking, the adaptation to mis-matched-room carrier will cause target identification to fall below the no-carrier baseline.

Another important issue is the duration of the preceding acoustic context needed for the perceptual system to get calibrated. For spatial hearing, there is evidence that localization performance in a weakly reverberant room can continue to improve after hours of exposure (Shinn-Cunningham, 2000). For speech perception, evidence from recent studies suggests more rapid adaptation timescales. Monaural compensation shows a continuous buildup of adaptation with increased exposure to a consistent previous context, at least up to 500 ms of exposure (Beeston et al., 2014). Intelligibility improves with longer exposure to consistent reverberation; the exposure duration at which performance asymptotes increases with SNR, from 850 ms for lower SNRs to 2.7s for higher SNRs (Brandewie and Zahorik, 2013). These results suggest that the buildup of adaptation to reverberation for speech perception occurs on a fairly rapid time scale. However, longer exposure to a preceding consistent environment may be more beneficial for more challenging listening environments without noise masking distorting the acoustic properties of the room. Furthermore, it is unclear whether additional exposure to an inconsistent carrier might be more detrimental for speech perception.

While past work explored the acoustic properties and the duration of the carrier, less emphasis has been given to non-acoustic factors, such as the ability to direct selective attention to the target speech. For example, knowing when target speech will appear might affect the ability both to benefit from a preceding consistent carrier and to overcome the disruption caused by an inconsistent carrier. Research on speech perception in complex auditory scenes suggests that prior knowledge of the spatial position and voice of a target speech can reduce attentional load and improve selective auditory attention and speech intelligibility in reverberation (e.g., Best et al., 2008; Shinn-Cunningham & Best, 2008). More research is needed to determine whether these top-down factors affect adaptation to reverberation.

Finally, it is not clear whether adaptation to reverberation generalizes across speech sounds and phonetic features with different acoustic properties. Adaptation does generalize across stimuli with diverse phonetic and lexical content, and, thus, is ecologically beneficial for real-world listening (Srinivashan and Zahorik, 2013). However, the use of lexical items does not enable a precise examination of adaptation processes at the segmental phoneme level, factoring out the contribution of higher order linguistic cues. A previously mentioned early study showed that monaural compensation mechanisms affect perception of the ‘sir’-‘stir’ contrast (Watkins, 2005). A later study showed that adaptation extends to more stops differing in place of articulation, especially /p/ and /b/ (Beeston et al., 2014). Stop consonants are popular candidates for studies investigating speech under adverse conditions, as they are particularly susceptible to masking by noise and temporal smearing by reverberation (e.g., Assman & Summerfield, 2004). Less is known about whether consistent room exposure improves the perception of other features that are also susceptible to room distortions (e.g., non-sibilant fricatives, place contrasts; Gelfand & Silman, 1979). A more detailed investigation of adaptation patterns across different speech sounds in different rooms can better inform theories and models of speech intelligibility in everyday listening environments.

Here, we performed two behavioral experiments that studied adaptation to room reverberation for consonant perception. In both experiments listeners were exposed to vowel-consonant (VC) syllables from a carrier phrase, followed by a target VC syllable simulated as being presented in one of two rooms, R1 or R2. The task was to identify the consonant from the final, target syllable. The carrier room was R1, R2, or anechoic space. The length of the carrier varied, containing either 2 or 4-VC syllables. Finally, the carrier/target uncertainty varied across the experiments. In Experiment 1 both the carrier length and the target room were randomly varied from trial to trial, i.e., participants could not predict when and from which simulated room the target would appear. In Experiment 2 the carrier length and the target room were fixed, i.e., participants knew in advance when and from which simulated room they would hear the target. Additionally, Experiment 1 also included no-carrier baseline trials, in which the target stimuli were presented in room R1, room R2, or an anechoic room without any preceding carrier.

The two reverberant rooms simulated in this study, R1 and R2, had broadband T_60_’s of approximately 3 s and 2.5 s, respectively. This strong reverberation was chosen to avoid performance ceiling effects that would preclude us from observing any benefits of adaptation. Previous studies have tackled this issue by using noise maskers (e.g., Zahorik and Brandewie, 2010). Although adding noise makes the task more difficult, the unique effects of reverberation and the listeners’ compensation mechanisms might differ. There is some evidence that adaptation to reverberation drops significantly at high levels of reverberation (Zahorik & Brandewie, 2016). We therefore expected differences in performance between the two rooms, and between the current and previous studies in which less reverberant rooms and noise maskers were typically used.

We tested these hypotheses in a series of analyses with the following structure. First, we examined the effects of consistent and inconsistent carriers relative to the baseline no-carrier condition of Exp. 1. Based on Brandewie & Zahorik (2018), we expected performance to be better for the same carrier, compared to no carrier. We examined whether the adaptation to an inconsistent carrier could cause performance to fall below baseline, an effect not observed previously. The duration of the 2 VC carrier exceeds the minimum duration leading to adaptation in past studies (e.g., 850 ms in Brandewie & Zahorik, 2016); we asked whether the longer, 4 VC carrier would further improve performance for the consistent carrier or further disrupt intelligibility for the inconsistent carrier.

Next, we contrasted performance across the two experiments to examine in detail how carrier length and target uncertainty affects speech intelligibility. Specifically, both the temporal position and room context of the target varied randomly between trials in Exp. 1 and were fixed in Exp. 2. Knowing the temporal configuration of the carrier and target syllables as well as the target room in advance might allow listeners to ignore the carrier, reducing attentional load and improving selective auditory attention to the target syllable. On the other hand, it is possible that if participants know when the target occurs, they may simply ignore the carrier and fail to calibrate to the carrier’s reverberation characteristics. This in turn is likely to reduce the effect of both consistent and inconsistent carriers. Such effects of temporal and contextual expectation across the two experiments are expected to interact with carrier room, and be greater for the longer carrier length.

Finally, an important goal of this study was to examine how consistent and inconsistent carriers affect performance across speech sounds with diverse spectrotemporal properties. To this end, in the last part of the analysis data were collapsed across experiments and carrier lengths, allowing us to examine effects of carrier room for individual phonemes and across their distinctive features of manner of articulation, place of articulation, and voicing.

Note that Exp. 1 was performed using a broader set of 16 consonants as stimuli. Since performance was at ceiling for 6 of those consonants, Exp. 2 presented only the remaining 10 consonants, although the participants could still use all 16 consonants when responding (see Methods section).

## Methods

### Participants

Nine young adult listeners participated in Experiment 1 and nine different adults in Experiment 2. All participants had normal hearing, as confirmed by an audiometric screening, and spoke English as their first language. All procedures were approved by the Boston University Institutional Review Board.

### Speech Material

Sixteen consonants (k, t, p, f, g, d, b, v, ð, m, n, ŋ, z, θ, s, and ∫) were used, each preceded by the vowel /a/. We used vowel-consonant (VC), rather than consonant-vowel syllables, as preliminary listening indicated that reverberation effects were greatest for final consonants (see also Gelfan & Silman, 1979). Stimuli were produced by three speakers, with one male recording taken from CUNY-NST corpus (Resnick et al., 1975) and one male and one female recording from the corpus described by Yund and Buckles (1995). For each VC, three tokens were spoken by each of three talkers. This resulted in a total of 144 unique speech tokens (16 VCs x 3 talkers x 3 tokens). Overall level differences across talkers were removed by equalizing the root-mean-square (RMS) energy levels of all tokens.

In Experiment 1 participants performed at ceiling for six of the consonants (k, t, n, z, s and ∫), with correct identification responses exceeding 90% in all tested conditions. Trials containing these consonants as target stimuli were removed from the analyses in Exp. 1 and these consonants were not included as targets in Exp. 2. However, in both experiments these stimuli were included in the carrier syllables and participants could still respond that they heard one of these consonants as targets.

For the feature-based analysis, the consonants were grouped by their manner of articulation, place of articulation, and voicing. Table 1 shows the feature classification used in this study.

**Table 1.**
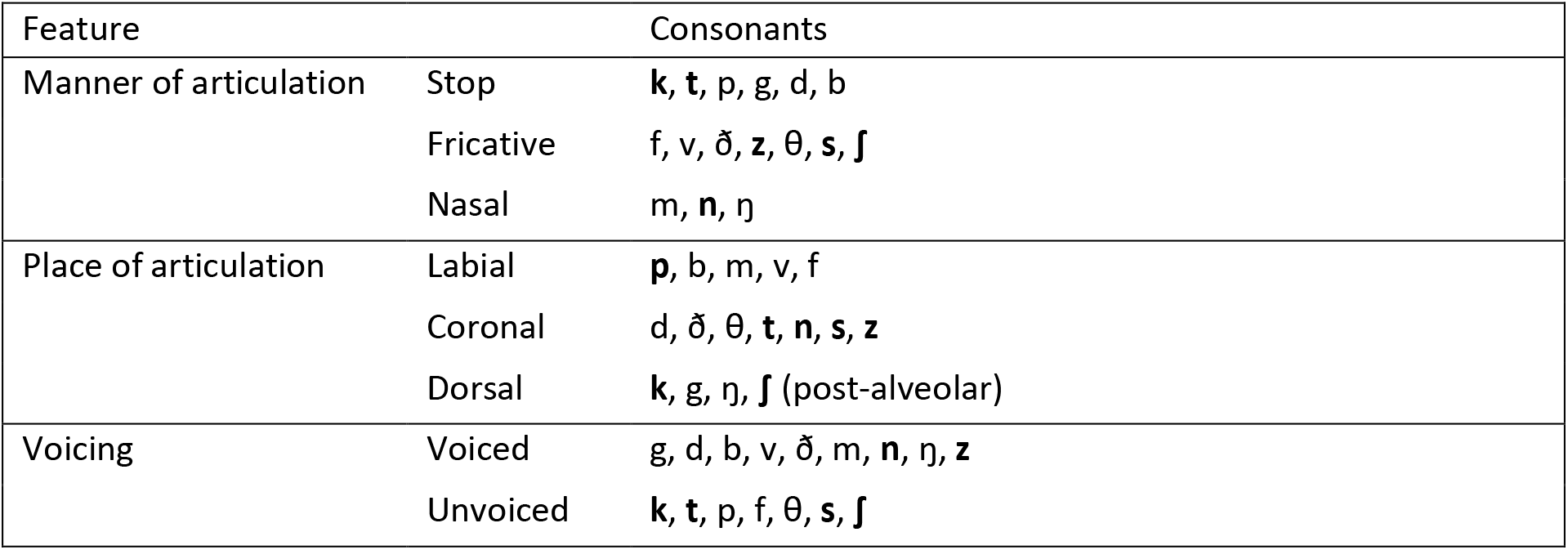
Phonetic feature classification. Consonants not used as target stimuli are in bold. All consonants were available as responses in both experiments.

### Room Simulation

To simulate the presentation of stimuli in different rooms, the VC tokens were convolved with binaural room impulse responses (BRIRs). The BRIRs were recorded using a setup consisting of an omni-directional (up to 2 kHz) dodecahedral loudspeaker system and a manikin head (Head Acoustics, HMM2) that faced the speaker system. BRIRs from two different large rooms were used, denoted as R1 and R2. The R1 response was measured in an elliptical church (room volume 13.333 m^3^) with the manikin relatively close (12 m) to the sound source, which was located beside the altar. The R2 response was measured in a large concert hall (room volume 22,776 m^3^, 2.020 seats) with the manikin located on the second balcony, 33 m from the speaker system located on the stage. The impulse responses were measured using the swept-sine method (Suzuki et al., 1995), for a time-stretched pulse of 1.35 seconds duration and with synchronous averaging (Satoh et al., 2004). An anechoic BRIR (AN) was derived from the R1 BRIR by time-windowing the first 5 ms of the response using a rectangular window to remove most of the reverberant energy. The resulting three BRIRs (R1, R2, and AN) were equalized for overall RMS energy. This equalization made the direct sound energy of R1, R2 and anechoic rooms quite different. However, the perceived loudness of speech stimuli convolved with the three BRIRs was comparable.

Figure 1 shows the acoustic properties of the BRIRs. Early time-domain portions of the responses in one ear are shown in Fig. 1A. R1 has a large echo around 50 ms after the direct sound, likely due to its elliptic room shape. Fig. 1B shows reverberation times (T_60_) as a function of frequency. R1 has a larger T_60_ than R2 at all frequencies. Fig. 1C shows the Clarity Index *C*_50_, i.e., the ratio of the early energy (0-50 ms) to the late energy (beyond 50 ms) in the impulse response as a function of frequency. *C*_50_ is lower in R1 than in R2, especially in the mid-frequency bands (250-1000 Hz). This analysis suggests that R1 should be more disruptive to speech perception than R2, while AN can be considered an ideal environment for speech perception, without any acoustic distortion.

**Figure 1.**
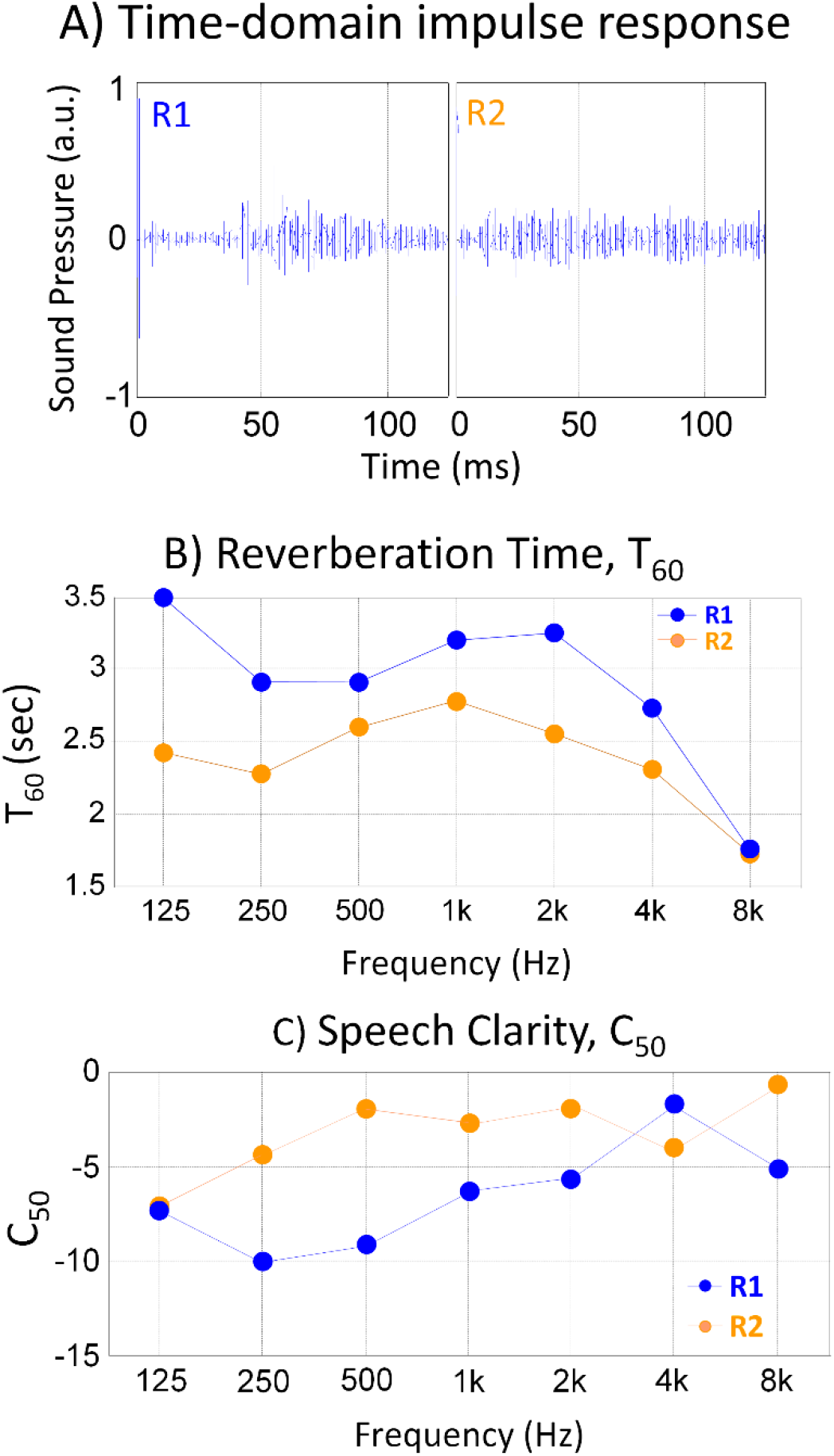
Acoustic properties of the BRIRs used in the Experiments. Blue and orange symbols are used for rooms R1 and R2, respectively. (A) Time-domain impulse responses from the left ear. (B) Reverberation time (T_60_), and (C), Clarity index (C_50_) as a function of frequency.

### Setup

In both experiments, performed in an experimental laboratory in the Boston University Hearing Research Center, participants were seated in front of an experimental computer inside a double-walled sound-proof booth. The experiments were implemented in MATLAB software (Mathworks Inc.). Stimuli were presented through a D/A converter (TDT RP2) and headphone amplifier (TDT HB7) driving insert headphones (Etymotic Research, ER1) at a comfortable listening level (adjusted by the experimenter). Participants responded using a graphical user interface (GUI) with 16 graphical buttons labeled with the 16 VCs, clicking with a computer mouse the button corresponding to the perceived target VC identity.

### Procedure

Prior to each experiment, a short training session was conducted to familiarize participants with the connection between the response GUI and the corresponding VC sounds. Participants were instructed to click on graphical buttons to produce the corresponding sounds, in a self-paced manner, until they felt confident about the relationship between sound and response. Upon clicking one of the buttons, a VC spoken by one male talker in an anechoic room was presented. There were no time constraints in this practice session, which typically took several minutes. Next, there was a short warm-up phase in which participants completed a session of 10 sample trials, identical to the test sessions described below. Participants were instructed to listen to the sounds and report the consonant in the final syllable. In Experiment 2 this warm-up phase was conducted each time the carrier length changed (described below).

The warm-up was followed by the experimental runs. On each experimental trial, participants heard an initial carrier, consisting of 2- or 4-VC syllables, followed by a target VC syllable. In Experiment 1 there was an additional control condition, in which participants only heard the target VC without a preceding carrier. The task was to report the consonant in the final target VC by mouse-clicking on the corresponding button in the GUI. The reverberation of the syllables in the carrier was randomly selected on each trial to be either R1, R2, or AN. With the exception of the no-carrier trials in Exp. 1, the reverberation of the target syllable was R1 on half the trials and R2 on the other half (in Exp. 1 no-carrier trials, R1, R2 and AN trials were presented with equal probability). The length of the preceding carrier (0, 2, or 4 VCs) varied randomly from trial to trial in Exp. 1, whereas it was blocked (2 or 4 VCs) in Exp. 2. Similarly, the target room varied randomly in Exp. 1 and was blocked in Exp. 2. The onset of each VC occurred 0.8 seconds after the onset of the previous VC. A random voice was selected for each trial and was consistent for all VC syllables within the trial. All three voices and three tokens per target VC were presented an equal number of times.

Each of the two experiments contained 720 trials in total. In Experiment 1 the trials were distributed across three sessions of 240 trials each. Each session contained (a) each of the 16 consonants in the target VC for each carrier length (2 and 4 VCs), carrier room (AN, R1, R2) and target reverberation (R1, R2) and (b) 48 control trials without a preceding carrier (No Carrier), with each of the 16 consonants as targets, for each room (R1, R2, AN). In Experiment 2, trials were distributed across two daily sessions of 360 trials. Each session contained two repetitions of each of the 10 consonants in the target VC for each of the three talkers and carrier reverberation (AN, R1, R2), in separate blocks for each carrier length (2 and 4 VCs). In each session the target reverberation was fixed (R1 or R2), with the order counterbalanced across participants.

### Statistical Analyses

For overall consonant identification, participants’ percent correct scores were logit transformed and entered into ANOVA tests. All figures show untransformed values and all error bars in the figures indicate SEMs. To quantify phonetic feature identification, we used the Information Transfer Rate (ITR) score, an information-theory derived measure (Shannon, 1948) commonly used for phonetic feature perception analyses (e.g., Miller & Niceley, 1955; Beeston et al., 2014; Sagi & Svirsky, 2008). The ITR is obtained by normalizing the mutual information between the speech stimuli and the participants’ responses by the stimulus entropy. A score of 1 indicates no confusions, whereas a score of 0 indicates random guessing. Unlike percent correct scores, this measure takes into account unbalanced categories and response biases (Sagi & Svirsky, 2008).

## Results

The results presentation is divided into two main parts. First, data are averaged across consonants, and the effects of carrier room, carrier length and carrier/target uncertainty are analyzed for the two target rooms. Then the data are combined across carrier length and carrier/target uncertainty and the effects of carrier room on the target rooms are analyzed separately for individual consonants and phonetic features.

### Data averaged across consonants

Figure 2 plots the across-participant-averaged percent correct responses from the two target rooms (shown by different color) as a function of the carrier rooms, for both experiments (rows) and all carrier-VC lengths (columns). Note that the no-carrier data were collected only in Exp. 1. (top left panel), and that this measurement was done also for the anechoic target room (pink). The horizontal lines extending from the top left panel mark the no-carrier baseline performance in the two reverberant target rooms, to allow a direct visualization of the effect of the different carrier rooms on performance.

**Figure 2.**
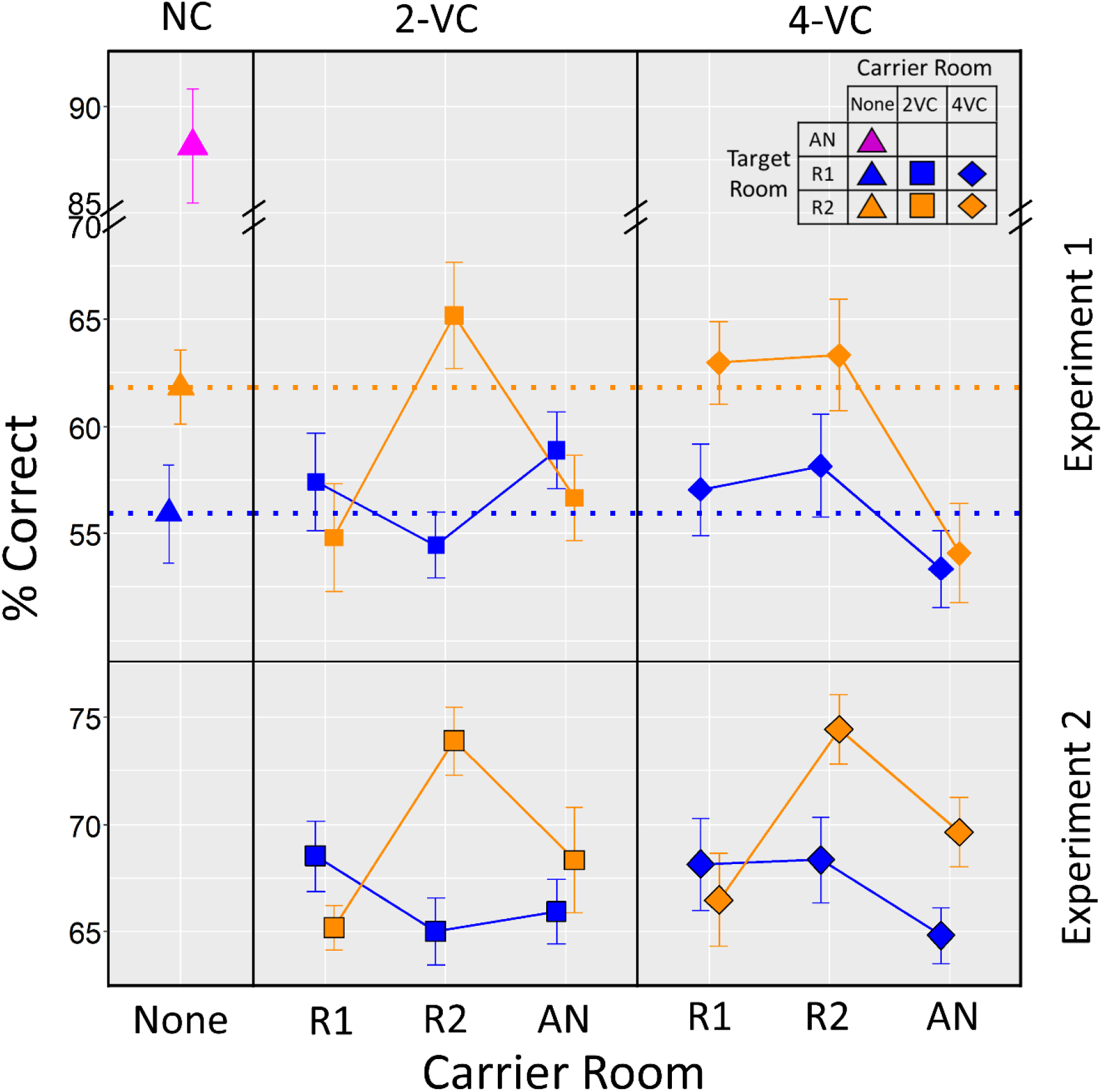
Across-participant average consonant identification accuracy (%) for Experiments 1 (top) and 2 (bottom) and for different carrier lengths (columns), plotted as a function of carrier room. Color represents target room. The no-carrier condition was only included in Exp. 1 (top left panel). The horizontal lines in the top panels show the no-carrier performance in the reverberant target rooms (orange and blue triangle from top left panel). Error bars show standard errors of the mean.

#### No-carrier baseline performance

The first analysis examined the participants’ baseline performance for targets not preceded by any carrier. The goals were to confirm that the reverberant rooms used in this study can degrade speech intelligibility significantly, and to establish a baseline for performance that would allow a direct evaluation of the effect of preceding carriers on consonant identification accuracy.

##### Results

The top left panel of Fig. 2 shows the mean identification accuracy in the baseline no-carrier condition of Exp. 1 for target stimuli simulated from all 3 rooms used in this study. The presence of reverberation had a dramatic effect on consonant intelligibility. While identification accuracy in the anechoic room reached almost 90%, in the two reverberant rooms it fell by about 30%. Further, the results show that intelligibility was higher for target room R2 (62%) than R1 (56%). Confirming these observations, a one-way repeated measures ANOVA with Target Room (R1, R2, anechoic) as the within-participants factor showed a significant effect (*F*(2,16) = 63.58, *p* = 0.0002).

##### Discussion

The comparison of no-carrier intelligibility in the anechoic room vs. the strongly reverberant rooms shows that the reverberation associated with the utterance of a vowel in a VC pair distorts perception of the subsequent consonant signal, interfering with identification. Note that performance degradation due to reverberation is likely to be much smaller for initial consonants (i.e., if the stimuli were CVs instead of VCs), as these consonants would not be affected by the vowel-related reverberation as much (e.g., Gelfand & Silman, 1979); specifically, the additional energy due to reverberation from a vowel will overlap the energy of a subsequent consonant, but not a preceding consonant. Informal piloting prior to the current study supported this prediction. It is also important to note that the masking effect of strong reverberation might be different from the masking effect of noise, which was used to limit the baseline performance in several previous studies performed in less reverberant rooms (e.g., Brandewie & Zahorik, 2018). Specifically, reverberation contains statistical regularities to which listeners are sensitive (Traer & McDermott, 2016). These regularities likely mitigate negative effects of reverberation in the current study, but not in studies with noise masking. Finally, the detrimental effect of reverberation was larger in room R1 than R2 in the current study. This is consistent with acoustic analysis showing a higher T_60_ and a lower C_50_ for this environment (see Fig. 1B-C).

#### Effect of a preceding carrier relative to the no-carrier baseline performance

The next goal was to examine the overall effect of a preceding carrier relative to the no-carrier baseline. Specifically, we tested whether an inconsistent carrier degrades performance relative to baseline and/or whether a consistent carrier would cause an improvement.

##### Results

The middle and right-hand upper panels of Fig. 2 show performance for target rooms R1 (blue symbols) and R2 (orange symbols) in the Exp. 1 conditions with a preceding carrier, while the color-matched horizontal lines show the no-carrier baseline performance (from the left-hand upper panel) for these rooms. Overall, the effect of carriers was much stronger for targets in room R2 than in R1 (compare the deviation of the orange symbols from the orange line vs. the deviation of the blue symbols from the blue line). A repeated-measures ANOVA with the factors of target room (R1, R2) and carrier room (No, R1-2VC, R2-2VC, Anechoic-2VC, R1-4VC, R2-4VC, Anechoic-4VC) showed an interaction between the two factors (*F*(6,48) = 2.58, *p* = 0.04), and significant main effects of each of them (target room: *F*(1,8) = 7.20, *p* = 0.0278; carrier room: *F*(6,48) = 3.13, *p* = 0.036). To assess whether there was a detrimental effect of inconsistent reverberation, a partial ANOVA was performed including only the no-carrier and the inconsistent-carrier rooms, with the factors of target room (R1, R2) and carrier room (No vs. average of Different-2VC, Different-4VC, Anechoic-2VC, Anechoic-4VC), where Different carrier refers to carrier from room R2 for target in room R1 and vice versa. This ANOVA found a significant main effect of carrier room (*F*(4,32) = 4.30, *p* = 0.0179; all other *p*’s > 0.15). Planned t-tests performed separately for each condition vs. the no-carrier in the same room found a significant degradation in performance induced by several of the conditions, with the strongest degradation caused by the anechoic 4-VC carrier for the R2 target room (drop in performance of 7.7%, *t* = 3.03, *p* = 0.016).

To assess whether there is a beneficial effect of consistent reverberation, a partial ANOVA was performed with the factors of target room (R1, R2) and carrier room (No, Same-2VC, Same-4VC), where Same carrier refers to R1 carrier for R1 targets and R2 carrier for R2 targets. This ANOVA found a significant main effect of target room (*F*(1,8) = 10.81, *p* = 0.011), while neither the main effect of the carrier nor the interaction of the two factors reached significance (*p*’s > 0.3). However, as can be seen in Fig. 2, there was a trend for performance to be better for the consistent carriers, in particular in R2.

##### Discussion

These results show only a weak trend of improvement in performance due to a consistent carrier, while previous studies showed that a consistent room carrier improves performance relative to a no carrier (e.g., Brandewie & Zahorik, 2018). However, we found that inconsistent room carriers can have a significant negative effect on consonant perception and even result in performance dropping below the no-carrier baseline. This result shows that tuning speech perception to the acoustics of one room can be detrimental when the subsequent speech is presented in a different room. This was particularly evident for target room R2, when the carrier was longer (4-VC) and presented in the anechoic room, and when the carrier was shorter (2-VC) and presented in either anechoic space or room R1. While it is not clear what factors determine that these are the most detrimental conditions, or why this detrimental effect was not observed in the previous studies, several aspects of the current study might be important. For example, it might be due to the very broad range of reverberation strengths imposed on carrier phrases, as the anechoic carrier is very dissimilar from the strongly reverberant rooms with T_60_’s of up to 3 s. Or, it might be that absence of additional masking noise from the stimuli in the current study allowed speech processing to tune more precisely to the acoustics of a particular simulated room, resulting in an increased negative impact when the room was switched. Such fine-tuning might not have been possible in the previous studies because identical noise was present in all conditions, masking some features of the presented speech. Finally, in the current study improvements and degradations relative to the no-carrier performance are stronger for the less reverberant target room R2, suggesting that there is a limit to the level of reverberation for which the auditory system can compensate to enhance speech perception, consistent with the previous reports showing the effects of preceding carriers diminish for rooms with very strong reverberation.

#### Effects of carrier room, carrier length, and carrier/target uncertainty

Three characteristics of the carriers and targets were systematically manipulated across the two experiments: the carrier room (same, different, anechoic), the carrier length (2 or 4 VCs), and the carrier/target uncertainty (in Exp. 1 the carrier length and the target room varied randomly from trial to trial, and thus listeners could not predict the target onset or its room; in Exp. 2 these parameters were fixed within a block). While there was no no-carrier condition in Exp. 2, the main prediction regarding the carrier room was that performance would be better after exposure to a consistent carrier, compared to either of the inconsistent carriers. Considering the two inconsistent carriers, Brandewie & Zahorik (2018) observed that a carrier with reverberation larger than the target was more disruptive than vice versa. Thus, a potential outcome was that the disruptive effect of the anechoic carrier would be smaller than that of either of the reverberant-room carriers. Alternatively, the anechoic carrier might be the most disruptive as the anechoic room was very dissimilar from both of the reverberant rooms, while the two reverberant rooms were relatively similar to each other. These effects of the carrier room were predicted to grow with carrier length, as it was expected that the tuning to each carrier room would get stronger over time, resulting in a larger improvement for the longer consistent carrier and a larger degradation for the longer inconsistent carriers. Regarding carrier/target uncertainty, it was expected that knowing when and from which room to expect the target might allow listeners to ignore the carrier altogether and focus attention exclusively on the target. This, in turn, would result in reduced interference from inconsistent carriers. Finally, while the two target rooms were both similar in that they were strongly reverberant, it was expected that the effects of carrier will be more visible in the less reverberant target room R2 than in R1, consistent with Zahorik & Brandewie (2016).

##### Results

To address these questions, the 2-VC and 4-VC data from Experiments 1 and 2 were analyzed together (all data from Fig. 2 except for the left-most panel), after recoding the carrier room factor levels as Same (carrier room reverberant, matching the target room), Different (carrier room reverberant, different from the target room), and Anechoic (carrier room anechoic, target room in R1 or R2). We ran a mixed ANOVA with carrier/target uncertainty (experiment) as a between-participants factor and with carrier room (Same, Different, Anechoic), carrier length (2 vs. 4 VCs) and target room (R1, R2) as within-participant factors. There were significant effects of carrier room (*F*(2,32) = 11.58, *p* = 0.0002), target room (*F*(1,16) = 6.30, *p* = 0.023) and experiment (*F*(1,16) = 5.54, *p* = 0.032), whereas carrier length had no significant main effect (*F* < 1, ns). There were also significant interactions of carrier length X carrier room (*F*(2,32) = 9.61, *p* = 0.0005) and carrier room X target room (*F*(2,32) = 8.11, *p* = 0.007). Finally, there was a significant triple interaction carrier room X carrier length X experiment (*F*(2,32) = 3.79, *p* = 0.033). All other interactions failed to reach significance (carrier room X target room X experiment: *F*(2,32) = 2.60, *p* = 0.117; in all other interactions *F*’s < 1, ns).

The strongest interaction found in the ANOVA was that of carrier room X carrier length, showing that there is a strong effect of the consistent-vs-inconsistent carriers on consonant identification, and that this effect is dependent on the carrier length. To better illustrate this interaction, Fig. 3A shows the data from Fig. 2 after collapsing across the two experiments and after rearranging the carrier room factor. This figure shows that the same-room carrier performance is independent of the carrier length (the leftmost square and diamond symbols are comparable for both target rooms). On the other hand, a significant effect of carrier length was observed with the different-room carrier, where performance was significantly more disrupted by the 2-VC carrier than the 4-VC carrier (compare the difference in performance between the middle square and diamond symbols). No effect of carrier length was observed for the anechoic carrier, similar to the same-room carrier. Statistical testing confirmed these observations. A partial ANOVA was performed that compared the effects of carrier room between the Same and Different carriers, including target room and carrier length as within-participant factors, and experiment as the between-participant factor. This ANOVA found a significant main effect of carrier room (*F*(1,16) = 15.31, *p* = 0.0012) and of target room (*F*(1,16) = 9.49, *p* = 0.0072). Importantly, there was a significant interaction of carrier room X carrier length (*F*(1,16) = 11.34, *p* = 0.0039), confirming that the disruptive effects of the different-room carrier depended on the carrier’s length. This interaction was independent of experiment and target room (interaction with experiment: *F*(1,16) = 3.21, *p* = 0.0923; interaction with target room: *F* < 1, ns; in all other higher order interactions *p*’s > 0.25). In contrast, and as expected, a partial ANOVA performed separately on the anechoic carrier data showed that carrier length did not affect performance (*F*(1,16) = 2.52, *p* = 0.1322).

**Figure 3.**
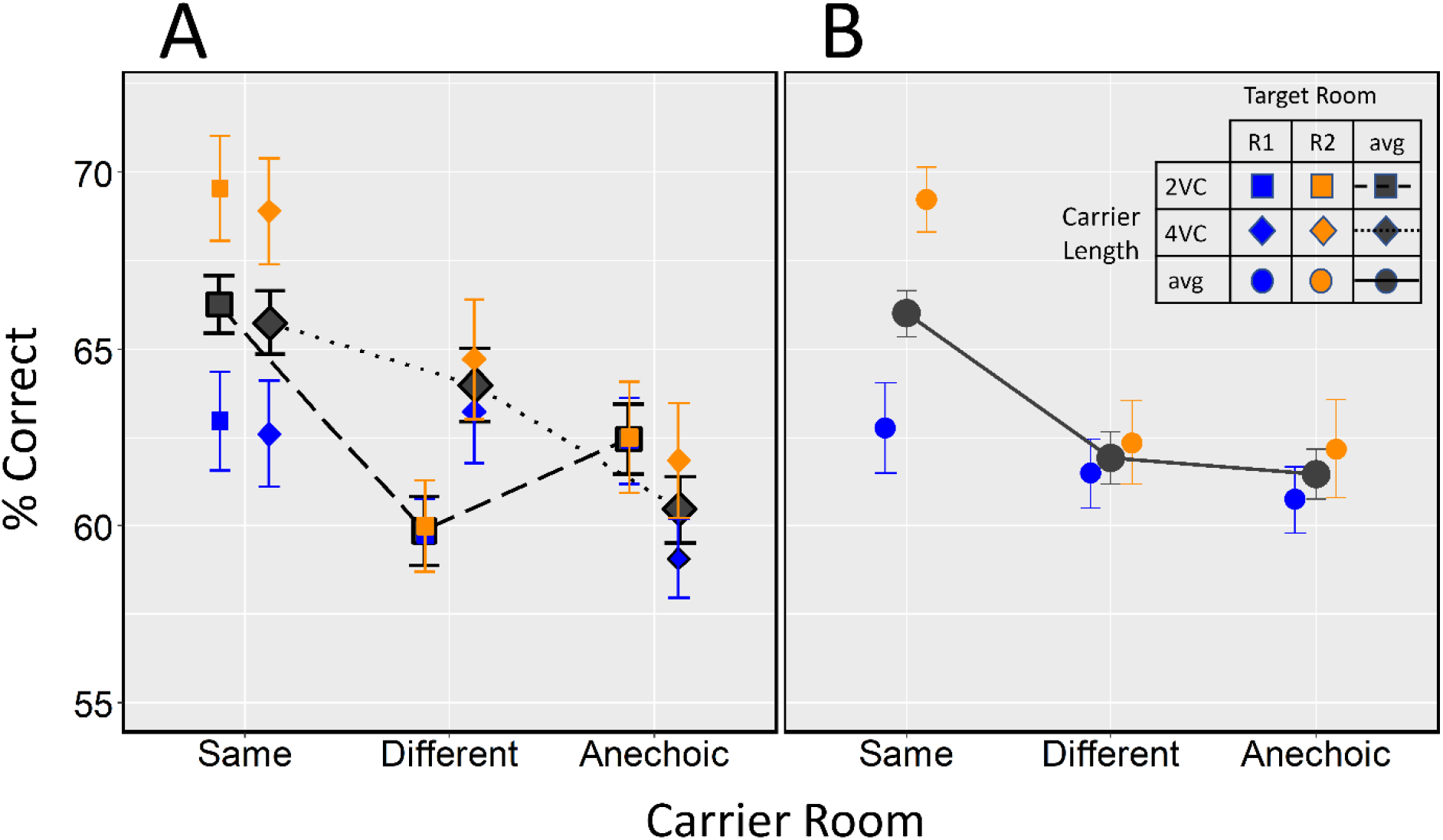
Mean consonant identification accuracy (% correct), collapsed across the two experiments as a function of carrier room. Data are plotted separately for target room R1 (blue) and R2 (orange), and as an average across target rooms (avg, black). (A) Square and diamond symbols show performance separately for the 2-VC and 4-VC carriers, respectively. (B) Circle symbols show performance averaged across carrier length. Error bars shows standard errors of the mean.

Finally, as before, the interaction carrier room X target room was significant (*F*(1,16) = 11.42, *p* = 0.0038). This result is better visualized in Fig. 3B, in which data are replotted after pooling across carrier length and experiment. To follow up on the significant interaction, we ran partial ANOVAs, separately for the Same data and for the Different and Anechoic data. The results show that the difference between the rooms is only present in the Same condition (*F*(1,17) = 16.59, *p* = 0.00079), while the Different and Anechoic conditions are very similar across the two target rooms (*F*’s <1, ns; blue and orange circles are far apart for the Same condition, but not for the Different or Anechoic conditions).

##### Discussion

The effect of carrier room, i.e., the magnitude of adaptation to reverberation, strongly interacted with the carrier length (Fig. 3A). It is unclear why, opposite to our expectations, the disruptive effects of the different-room carrier were stronger in the shorter than the longer carrier, and why this interaction is not observed for the anechoic carrier. A potential explanation is that the short carrier is more disruptive because both the short distractor and the target are automatically processed as one object, perhaps because cognitive load connected with processing the short stimulus is large, while for the longer distractor the participants are able to switch from the carrier to the target, as if they are two objects, and to treat the target as a new object for which the reverberation characteristics are estimated separately. This might also explain why the disruptive effects of the anechoic carrier were independent of carrier length, as the anechoic room is drastically different from the reverberant targets, making it easier to separate the carriers from targets and treat them as distinct objects, regardless of the length of the carrier.

The carrier room x carrier length interaction was also weakly dependent on the experiment factor, i.e., on carrier/target uncertainty, as evidenced by the significant 3-way interaction. However, this weak interaction did not come out as significant when partial ANOVAs were performed. Also, it was expected that the effect of uncertainty would be evident for 2 VCs because that was the only length for which the participants could not know ahead of time whether they are hearing 2-VCs or 4-VCs, as in a 4-VC trial the participant could realize the carrier had 4-VCs simply from the fact that a 4^th^ VC was presented. Instead, an effect of carrier/target uncertainty was observed for 4-VC and only in one room (R1-carrier R2-target in Fig. 2). Therefore, this weak interaction was not considered further.

The effect of carrier room also varied with target room, as reflected in the significant carrier room X target room interaction (Fig. 3B); this result is largely consistent with results when the data are analyzed separately for different VC-lengths, as shown in Fig. 3A. Specifically, results from the 2 target rooms only differ in the Same condition. However, Exp. 1 results suggest that the main effect of the carrier is a disruptive effect from the inconsistent carriers, not a beneficial effect of a consistent carrier. Thus, the difference in the Same performance between the rooms is likely due to the difference in baseline performance in the two rooms, and to the fact that the disruptive effect of the inconsistent carriers (Different and Anechoic) is much larger for the target room R2 than for R1.

In sum, the results of analysis on data averaged across consonants suggest that the effect of carrier room is caused mainly by perceptual disruption of an inconsistent carrier, rather than a beneficial effect of the consistent carrier. The effects of carrier length show up as a significant interaction for the different-room carrier, with performance worse for the short carrier. The magnitude of the effect is room dependent and is smaller in the more challenging reverberant target environment. This finding partially confirms past results, showing greater disruption for a less reverberant target room (Zahorik & Brandewie, 2016). However, the anechoic carrier was as disruptive as the different carrier. Therefore, it appears that the overall acoustic dissimilarity of the carrier and target rooms also plays an important role. In contrast to the factors of carrier room, target room, and carrier length, there was very little evidence that prior knowledge of the target onset and room (i.e., the carrier/target uncertainty) affected performance.

### Adaptation to reverberation for individual consonants and phonetic features

Two analyses were performed on data pooled across the factors of carrier length and carrier/target uncertainty (i.e., corresponding to the across-consonant analysis in Fig. 3B) to examine how perception of individual phonemes and of phonetic features is affected by a previous exposure to speech from a consistent or inconsistent room.

#### Results

Fig. 4 shows the listeners’ perceptual confusions separately for each carrier room (left-hand vs. center vs. right-hand panels) and target room (upper vs. lower panels). Each cell [*i,j*] within a panel shows the across-participant average percentage of times that the stimulus in column *j* was identified as the response in row *i*. To improve legibility of the tables, the cells containing the value of 0 were left blank, and different shades of red were used in the background of each cell to match the numeric value in it (darker red corresponding to a higher value). Note that participants were not limited to the stimulus set in their responses (they could respond with any one of the 16 consonants described in Table 1), and thus confusion matrices are not square. Therefore, some of the cells that correspond to correct responses, highlighted by a blue frame, are slightly off the primary diagonal. The consonants in the confusion matrices are grouped hierarchically by voice, place, and manner of articulation, as indicated by labeling along the x-axis and y-axis of the graph.

**Figure 4.**
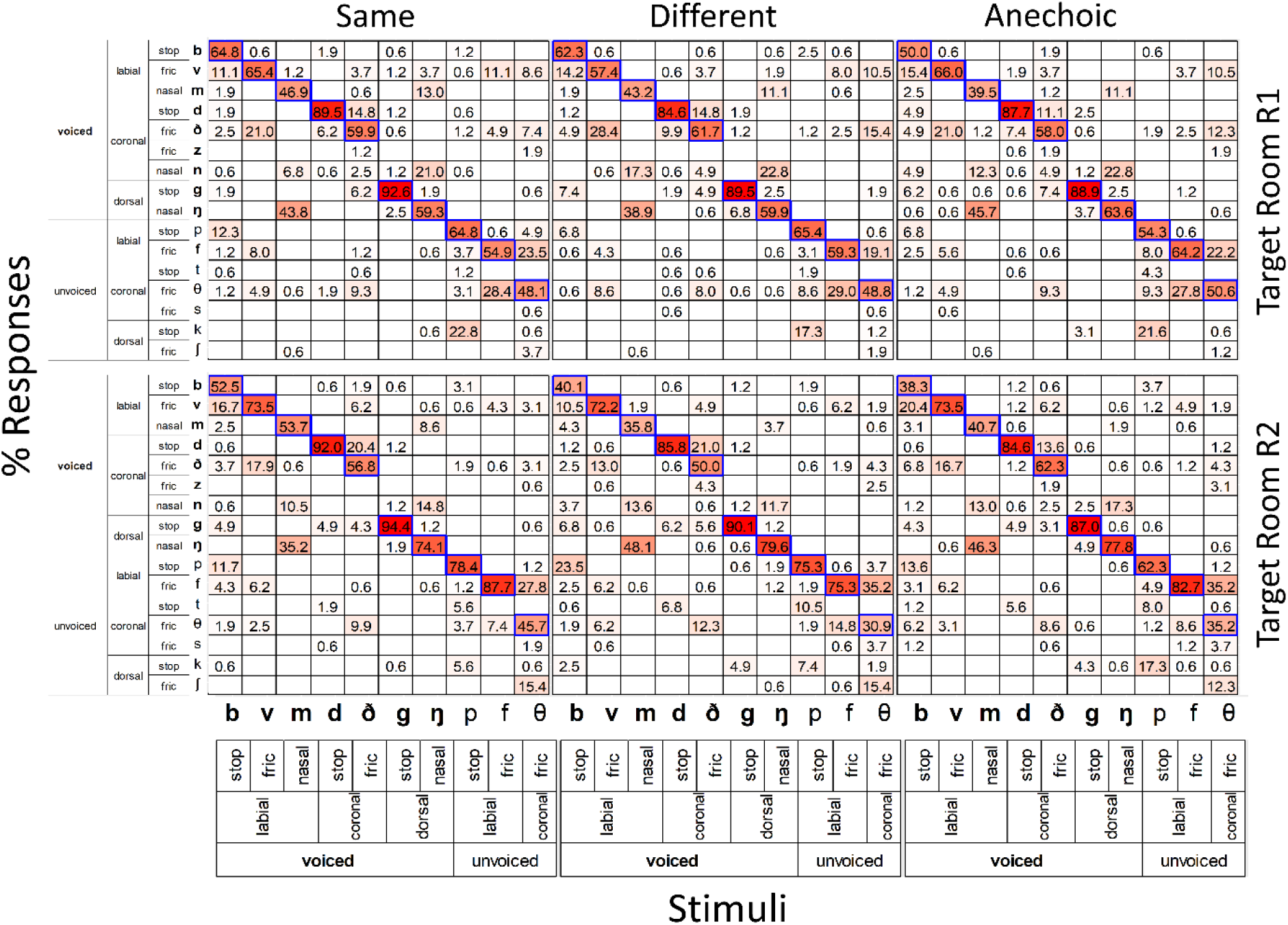
Consonant confusion matrices. Across-participant-average confusion matrices, pooled across the two experiments and carrier lengths. Separate matrices are shown for the Same (left-most panel), Different (middle) and Anechoic (right-most panel) carriers, and for each Target Room (R1, top; R2 bottom). Columns show the actual speech stimuli that were presented and rows show the response options. Each cell i,j shows the percentage of times the consonant in column j was identified as the consonant in row i (empty cells denote 0 percentage). White colors in the tiles represent lower and red colors higher stimulus-response percentage. The legends shown along the vertical and horizontal axes denote consonant classification according to voicing (bold letters for voiced and plain for unvoiced), manner of articulation and place of articulation. Blue frame highlights the cells that represent correct responses.

As can be seen from Fig. 4, performance varied considerably across phonemes. The two consonants more severely affected by reverberation were /m/ and /θ/, with overall identification accuracy less than 45%. At the other extreme, /g/ and /d/ were perceived much more accurately, with average performance exceeding 85% correct. A closer examination of the participants’ errors reveals that, for each stimulus, confusions clustered around one or two dominant responses that tended to be consistent across carrier and target rooms. Specifically, for 8 out of the 10 target consonants, the primary confusion was consistent across all, or most, carrier and target rooms (6/6 conditions for 5 consonants /v→ð, m→ŋ, ð→d, f→ð, ð→ f/, and 5/6 conditions for 3 consonants /g→ŋ, ŋ→n, p→k/) and accounted, on average, for 52 % of the total error. Further examination of the participants’ errors revealed that a few phonemes were mutually confusable, with the clearest cases being /θ-f/ (for both target rooms) and /d-ð/ (for the R1 target room). However, in most cases the phonemes were not equally confusable with each other. Specifically, even though nasals were mainly confused with each other, e.g., /m/ was systematically confused with /ŋ/, for /ŋ/ the primary confusion was /n/ (which remained a response option even when not presented as a target consonant) and /m/ was the secondary confusion. For /b/ the primary confusion was /v/ (in 4/6 conditions), whereas /v/ was consistently confused with /ð/. This and other examples suggest that reverberation created consistent consonant confusion groups that were mostly asymmetrical. Finally, an examination of the responses at the feature level showed that for each stimulus, the more frequent responses involved the correct detection of at least 2 features. There was only one instance in which participants’ missed all 3 features more than 5%. The phoneme /b/ was perceived as /θ/ in the anechoic carrier room and target room R2. Importantly, when considering these observations, note that across-participant differences were strong (not shown).

To identify adaptation-to-reverberation processes at the phonetic feature level, the analysis next focused of the effect of a preceding carrier on place of articulation, manner of articulation, and voicing. Specifically, after grouping the consonants into different categories according to these features (see Table 1 and the category labels along the x-axis and y-axis of Fig. 4), new confusion matrices were derived from those in Fig. 4, separately for place, manner, and voicing. In these confusion matrices, the stimulus response pairs were only considered at the feature level, i.e., identifying the voiced labial stop of /b/ as an unvoiced labial stop of /p/ would increase the number of voiced-unvoiced (i.e., incorrect) responses in the voicing feature category, labial-labial (i.e., correct) responses in the place category, and stop-stop (i.e., correct) responses in the manner category. Based on these new matrices, we computed for each individual participant the information transfer rate (ITR) for each feature across the different combinations of carrier and target rooms.

Figure 5 shows the across-participant average ITR score as a function of carrier room, separately for each phonetic feature (separate panels) and each target reverberation (different colors within each panel). Consistent with our previous results, overall performance tended to be higher for target room R2 (orange) and for the same carrier condition. To evaluate the effect of carrier on each of the phonetic features, a separate two-way repeated measure ANOVA with the factors of carrier room and target room was performed on ITR values for each of the features. For manner, this ANOVA yielded a significant main effect of carrier room (*F*(2,34) = 4.03, *p* = 0.027) and target room (*F*(1,17) = 5.73, *p* = 0.029), while the interaction was not significant (*p* > 0.3). For place, a significant main effect of target room was found (*F*(1,17) = 13.82, *p* = 0.002), while neither the main effect of carrier nor the interaction were significant (*p* > 0.2). For voicing, only the interaction of the two factors was significant (*F*(2,34) = 5.78, *p* = 0.007), while the main effects were not significant (*p* > 0.1). Post-hoc planned comparison performed as a paired t-test on the contrast of consistent vs. inconsistent (Same – 0.5*[Different+Anechoic]), separately for each target room and each phonetic feature, found a significant difference (*p* < 0.5) for manner in target room R1 and for voicing in target room R2 (see horizontal lines and asterisks in Fig. 5).

**Fig. 5.**
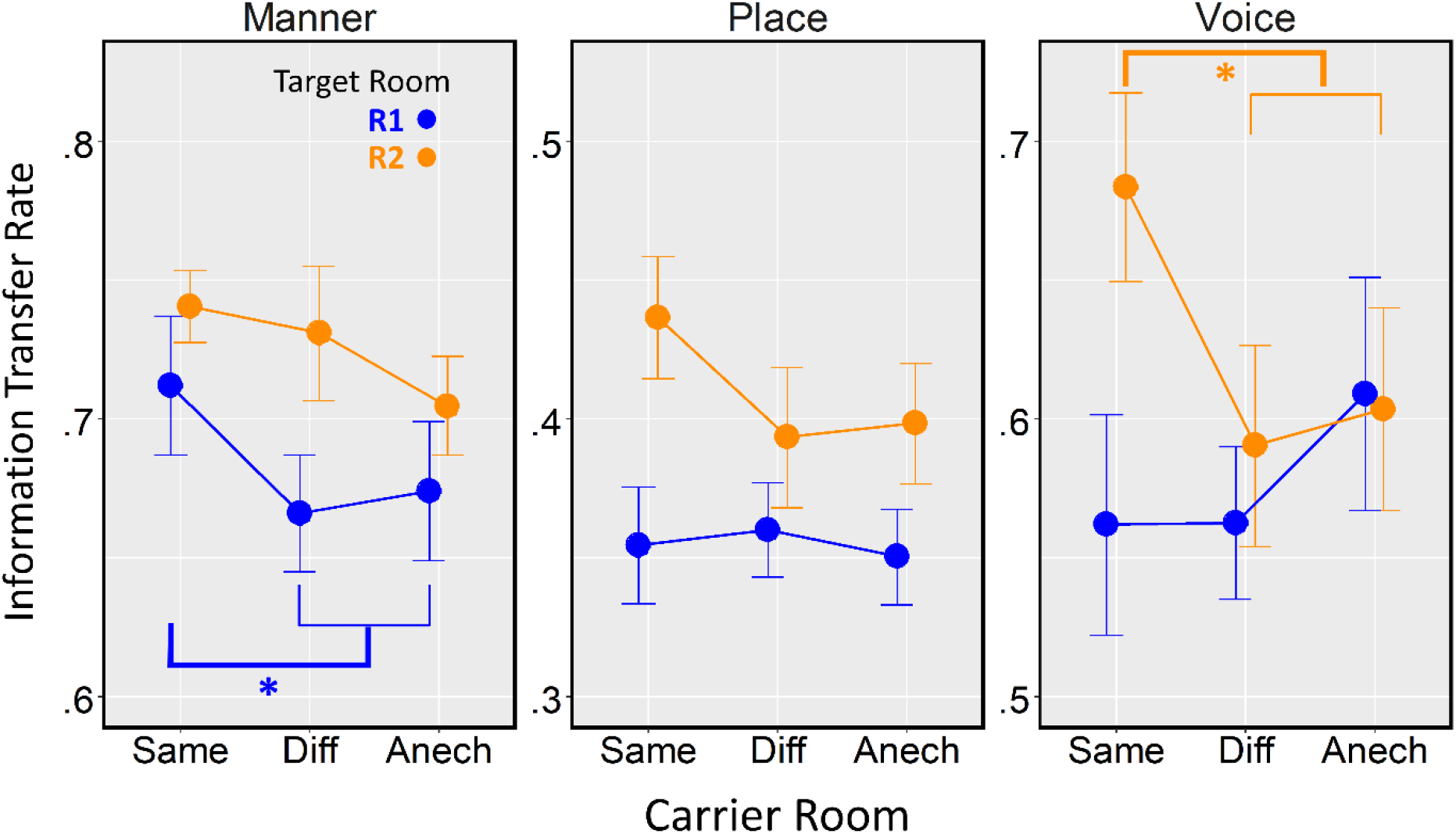
Across-participant average Information Transfer Rate (ITR) as a function of carrier room for manner or articulation, place of articulation and voicing, separately for target room R1 and R2.

To further examine the significant ITR improvements with consistent vs. inconsistent carriers, Fig. 6 plots the percent correct identification of individual phonetic features corresponding to manner of articulation (left-hand panel, showing both target rooms) and voice (right-hand panel, only for target room R2). The left-hand panel shows that the benefit of consistent carrier affects only the stop consonants, while the right-hand panel shows that the voice ITR improvement is caused mainly by improvement in identification of the voiced consonants. Confirming these observations, paired t-tests showed significant improvements for stop consonants (*t(17)* = 3.58, *p* = 0.0023) and voiced consonants (*t(17)* = 3.48, *p* = 0.0029), while for the unvoiced consonants the difference was not significant (*p* > 0.5).

**Fig. 6.**
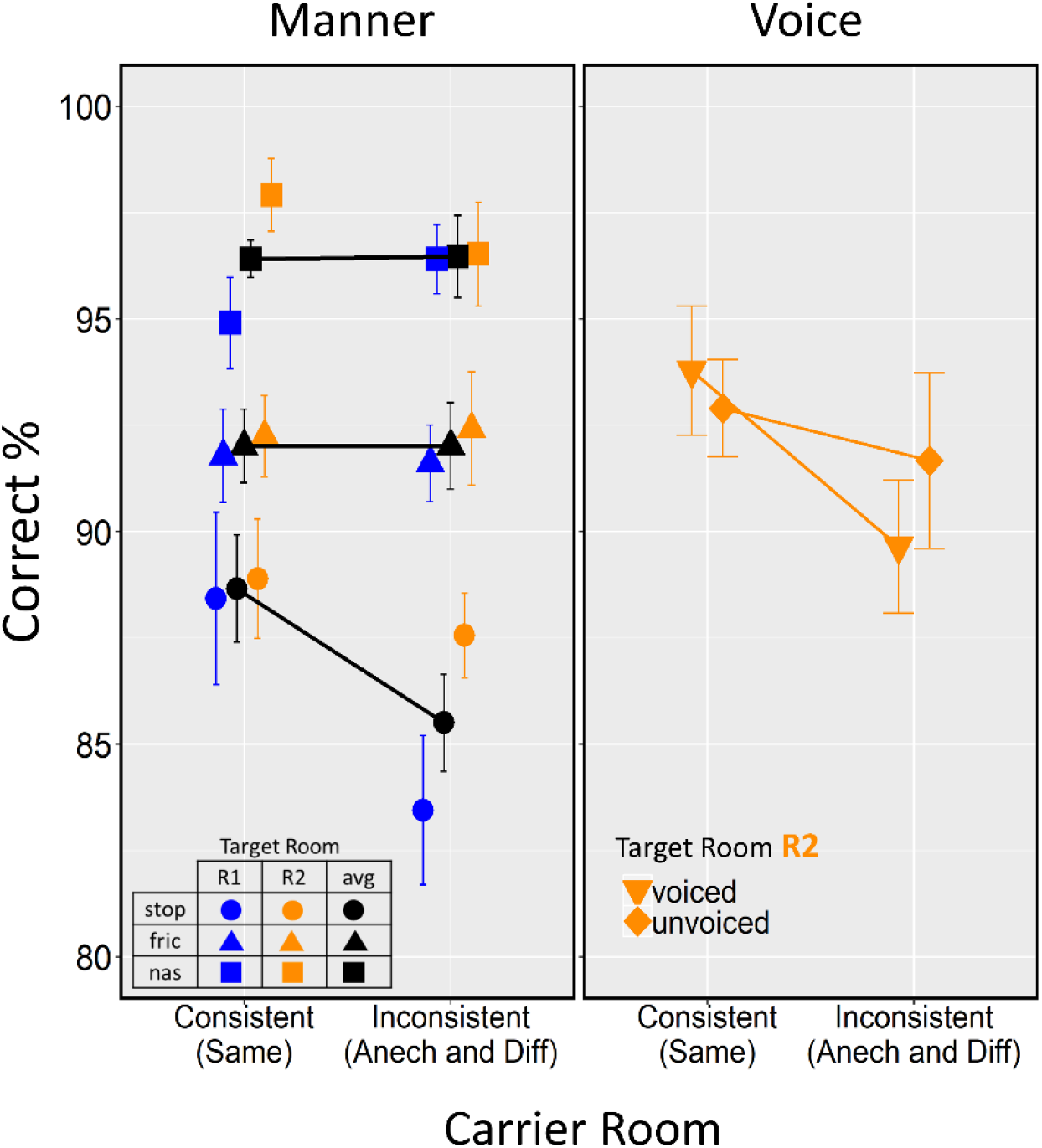
Across-participant average percent correct feature identification as a function of carrier room. Symbols denote the individual phonetic features. Manner or articulation data (left-hand panel) are plotted separately for target room R1 (blue) and R2 (orange), and as an average across target rooms (avg, black). Voicing data (right-hand panel) show performance for target room R2. Error bars show SEMs.

#### Discussion

Fig. 5 shows that, for both rooms, manner of articulation was the feature with the highest transmission (ITRs ranging from .65-.75), followed by voicing (ITRs from .55-.7) and place of articulation (ITRs from .35-.45). The particularly low performance for place is consistent with previous work on phonetic confusions in noise and reverberation, showing that place is negatively affected, especially for consonants in the final position (e.g., Gelfand & Silman, 1979; Miller & Nicely, 1955).

Manner of articulation (left-most panel in Fig. 5) is the feature for which a consistent carrier yields the greatest benefit compared to a carrier from an inconsistent room. This improvement was observed in both rooms for the stop consonants, whereas for fricatives and nasals there was no evidence of improvement (Fig. 6). On the other hand, no improvement for the same-vs-different carrier was observed for the place feature (central panel in Fig. 5), even though there was a trend for same-carrier improvement in target Room R2. Finally, the feature of voicing (right-most panel in Fig. 5) showed a strong same-vs-different carrier improvement for target room R2, but no such effect for R1. This improvement was driven primarily by the voiced consonants (Fig. 6). The room specificity of this effect suggests that tuning to the voicing characteristics in the more reverberant R1 carrier has a negative effect on consonant identification in the less reverberant R2 target, but not vice versa. Overall, these results confirm previous reports that stop consonants are affected the most by adaptation to room reverberation. They also show that voiced consonants can be affected by adaptation in certain rooms.

## General Discussion

This study investigated how consonant perception in a highly reverberant room is influenced by a preceding carrier phrase simulated from either the same or a different room. The effects of various combinations of carrier and target rooms were examined using natural reverberation, without adding noise or introducing other manipulations, such as abrupt cut-offs, that have been used in previous work (e.g., Brandewie & Zahorik, 2018; Zahorik & Brandewie, 2016; Srinivasan & Zahorik, 2013; Beeston et al., 2014). Here, for two reverberant target rooms, we examined different aspects of the preceding carrier and target: the carrier room (i.e., the preceding carrier either had the same room reverberation as the target, a different room reverberation, or was anechoic), the carrier length (either 2 or 4-VC syllables), and the carrier/target uncertainty (the carrier length and target room were either fixed or varied randomly from trial to trial). The results were analyzed either averaged across the examined consonants or after grouping consonants along three phonetic features (manner of articulation, place of articulation, and voicing).

Without a preceding carrier the more reverberant of the two simulated rooms degraded perception of some consonants, but had a negligible effect on others. Specifically, the target consonants /z, n, t, s, k and ∫/ included in Exp. 1 were removed from further analysis because their perception was largely unaffected by the room reverberation. Not surprisingly, 3 of these 6 consonants were sibilants, with strong energy at higher frequencies, which have been shown to be resistant to both noise and reverberation (e.g., Gelfand & Silman, 1979; Danhauer & Johnson, 1991; Miller & Nicelly, 1955). Performance was also unaffected by the room acoustics for the unvoiced stop consonants /t/ and /k/, while it significantly dropped for the unvoiced stop /p/. While it is outside the scope of this study to determine why reverberation affects some consonants more than others, it is possible that the strong high- and mid-frequency bursts, respectively, that are critical for the perception of /k/ and /t/ survived reverberation, in contrast to /p/, which is instead characterized by a soft wide-band click that diminishes to a low frequency burst (Li & Allen, 2011; Li et al., 2010), making it more susceptible to smearing by reverberation. Overall, in agreement with previous studies our results show that there is considerable variability in how reverberation affects different speech sounds, ranging from negligible, to moderate, to strong disruptions in perception (e.g., Danhauer & Johnson, 1991; Gelfand & Silman, 1979).

Averaged across the remaining 10 consonants, in Exp. 1 we expected to find a significant improvement in speech perception after exposure to a consistent carrier, relative to a no-carrier baseline condition (e.g., Brandewie & Zahorik, 2010; Beeston et al., 2014; Srinivashan & Zahorik, 2013, etc.). However, we only found a weak improvement in overall identification accuracy. On the other hand, the inconsistent carriers, on average, impaired performance compared to the no-carrier baseline. Thus, overall, the negative effect of inconsistent carriers *re*. baseline was stronger than the positive effect of consistent carriers, while in previous reports an inconsistent carrier never led to worse performance than the no-carrier baseline (Brandewie & Zahorik, 2018). This result suggests that in strongly reverberant environments in which the effect of carrier adaptation can be measured even without noise masking, listeners are less able to take advantage of a consistent preceding context to improve perception, while, at the same time, they are very susceptible to the disruptive effects of an inconsistent context — either anechoic or in different reverberation.

The remaining analyses considered data from both Exps. 1 and 2, collected with carrier length and target room random or fixed, respectively. First, the effects of carrier room and carrier length on across-consonant average performance were examined. Depending on the carrier length, the duration of the preceding carrier was either 1.6 s (2-VC carrier) or 3.2 s (4-VC carrier). Our results suggest that, for such brief carriers, exposure to the longer consistent carrier did not improve performance more than the shorter consistent carrier. This is consistent with our previous analysis on the Exp. 1 data, where we did not observe significant improvement in the consistent carrier relative to the no carrier baseline condition. It is also in line with previous reports which suggest that, at low levels of noise masking, such buildup reaches its maximum within 850 ms (Brandewie & Zahorik, 2013).

On the other hand, for inconsistent carriers, surprisingly, the effect of carrier length was non-monotonic: for the mismatched reverberant room, performance was worse with the 2-VC compared to the 4-VC different-room carrier. No such interaction was observed for the anechoic carrier. It is unclear why this non-monotonicity was observed. If anything, we expected the disruptive effect of the longer inconsistent carrier to build up over time and to be stronger than that of the shorter carrier. One possible explanation is that the short carrier is more disruptive because it is automatically processed as one object with the target. The longer distracting carrier, on the other hand, might help listeners to reset auditory streaming and treat the carrier and the target as distinct objects. Similar effects were observed in streaming/grouping studies of sound localization (Kopco et al., 2017) and, to some extent, also in speech perception in multi-talker environments (Best et al., 2008). This might also explain why carrier length did not influence how the anechoic carrier impacted performance. Specifically, given that the anechoic room is dramatically different from either of the reverberant rooms examined here, it is likely that the anechoic carriers were always perceived as separate streams from the reverberant targets. However, this explanation needs to be tested in the future. In any case, our results suggest that, rather than being more disrupted by longer exposure to an inconsistent carrier, listeners might actually be able to exploit its longer duration to overcome detrimental effects of reverberant energy on speech perception.

The two different types of inconsistent carriers used in this study were expected to affect performance differently. On the one hand, the anechoic carrier might be more disruptive than the different-room reverberant carrier, as it has substantially different acoustic characteristics than both reverberant target rooms. On the other hand, the anechoic carrier does not distort the stimuli, giving the listeners a chance to have a good “look” at the “clean” version of each phoneme. Such looks may be beneficial when identifying target speech distorted by reverberant energy. Contrary to our predictions our results showed that, averaged across carrier lengths, there was a similar drop in performance for the anechoic and different-room reverberant carriers, perhaps because the two above-mentioned contradicting effects of the anechoic carrier tend to cancel.

Our results failed to show a clear effect of carrier/target uncertainty on the across-phoneme averaged data. This is in line with previous reports that uncertainty about the temporal location of the target stimulus does not reduce the magnitude of adaptation to reverberation (Beeston et al., 2014). However, due to the relatively small sample size in our study, firm conclusions should be avoided and more research is needed to examine the effects of directing selective attention to the target speech in reverberation.

A persistent finding in our study was that the effect of the different carrier rooms was much smaller for the targets in room R1, as reflected in the significant interaction between carrier room and target room. This is likely due to the larger broadband T_60_ of R1, which resulted in a marked decrease in performance in the no-carrier condition. This strong extra reverberation not only made the baseline performance worse, it also made it more difficult for listeners to benefit from prior exposure to this room. There was also only a little negative effect of inconsistent carriers on R1 targets, consistent with a previous report that the more reverberant inconsistent carriers have a more negative effect on the less reverberant target than vice versa (Brandewie and Zahorik, 2016). An alternative explanation is that R1 had a more detrimental effect on consonant perception than R2 not because of its stronger reverberation but due to its peculiar characteristics, such as its elliptical shape and prominent low frequency resonances. It is important to address this second possibility by controlling the level of reverberation while using more rooms, with different wall materials and layouts. Consistent with both accounts, our results suggest that the magnitude of facilitation or disruption due to adaptation to reverberation can vary considerably depending on the acoustic properties of the target room.

The second major goal of this study was to examine whether the relative benefit of consistent vs. inconsistent carrier phrases for consonant perception was specific to certain phonetic features, e.g., the previously reported stop consonants, or whether it also affects other features that are representative of the acoustic-phonetic diversity of everyday listening. An examination of confusions for each phoneme revealed that different carrier rooms have different effects on performance. Phonetic feature analysis showed that the highest information transmission was observed for manner of articulation, followed by voicing and place of articulation. This is consistent with previous work on phonetic confusions in noise and reverberation (e.g., Miller & Nicely, 1955; Gelfand & Silman, 1979). The larger number of place errors is also consistent with previous reports (e.g., Miller & Nicely, 1955; Benki, 2004). Manner was the feature that showed the most robust improvement in performance for the same-room carrier, which was observed in both reverberant target rooms and was restricted to the stop consonants. For voicing there was a large consistent-vs-inconsistent carrier difference for the R2 target room, but no difference for the more reverberant R1 target room. This asymmetry might help explain the above-mentioned asymmetry in how much degradation was caused by the inconsistent-room carrier for target room R1 vs. R2 in the across-consonant average data. Also, it might be the cause of the previous report that there is a greater disruption in speech identification caused by a more reverberant carrier than by a less reverberant carrier (Brandewie & Zahorik, 2016). Specifically, the current results suggest that this asymmetry is driven primarily by specific disruptions in the identification of voicing, and specifically for the voiced consonants, for which the detrimental effect was significant. Finally, for place of articulation there was only a weak trend for an improved performance on the same carrier in the R2 target room that did not reach significance. Thus, the place of articulation seems to be the feature that is the least affected by the specific characteristics of any given reverberant room and/or the characteristic to which the auditory system is tuning the least when adapting to a specific reverberant room.

This study has some limitations. First, the two rooms used here, while natural and realistic, have higher levels of reverberation than environments in which typical listeners spend the majority of their daily lives. This choice was motivated by our goal of directly examining the effects of reverberation on consonant perception by testing difficult conditions without combining it with the effect of noise masking. However, as has been shown by previous research, the benefit of a consistent carrier is diminished in very strongly reverberant target rooms (Brandewie & Zahorik, 2016). This was the case in our study, where very little improvement in consistent-carrier performance was observed even after removing six of the original consonants that were unaffected by room reverberation. On the other hand, a higher baseline means that the disruptive effects of inconsistent carriers are more likely to be visible, as we find in the current study. Therefore, it should be noted that our results are likely to generalize to challenging environments such as churches, large lecture halls or concerts halls, but may not explain effects in modestly reverberant environments. Second, although we tested a number of phonetic units and we examined three carrier rooms, we included only two target rooms, with particular acoustic characteristics. Future studies should include additional strongly reverberant environments with different geometry and reverberation time. Similarly, the current study only analyzed 10 consonants preceded by a single vowel. While beyond the scope of this study, we believe that these different sources of variability need to be addressed in future studies in order to obtain more generalizable findings for adaptation of speech perception to reverberation. Finally, the reverberant tails of the stimuli in the current study extended beyond the gap between individual VCs in each stimulus. Thus, theoretically, the reverberant carrier VCs might have energetically affected the target VCs. Previous studies artificially removed a portion of the carrier reverberant tail to avoid any artifacts caused by the overlap of reverberation from a preceding VC during target VC presentation. Here, no such modifications were made, as the signal during the presentation of the consonant in the target VC was dominated by the immediately preceding vowel and its reverberation, and thus the energetic effect of the carrier VC tails was minimal.

In sum, the current results partially confirm the results of previous work while, at the same time, point to a more complicated picture for consonant perception in reverberation (Helfer, 1994). We found that for consonants presented in particularly challenging rooms without masking noise, the effects of a preceding acoustic context on speech perception manifest as a disruption by inconsistent carriers, rather than an improvement by a consistent carrier. In addition, listeners seem to be able to exploit longer exposure to an inconsistent carrier in order to process the carrier and targets separately and overcome the detrimental effects of the inconsistent carrier. These effects of the preceding carrier are less robust for a more reverberant target room, suggesting that there is an upper limit to the degree of reverberation to which the auditory system can compensate. Finally, the effects of the preceding carrier affect certain phonemes and phonetic features more than others. Performance for manner of articulation and, partially, for voicing is improved after exposure to a consistent relative to an inconsistent carrier, while place of articulation is not affected. Although previous research has revealed important insights about adaptation to reverberation for speech perception, to our knowledge, this study is the first to show the patterns of improvement and disruption for high level of reverberation without masking, while examining a large set of consonants that represent much of a language’s phonetic repertoire. Clearly more research is needed to determine how listeners are able to overcome the disruptive effects of inconsistent carriers to understand speech in very challenging listening environments and when moving from one environment to another. Such understanding might also be useful for the development of prosthetic devices for the hearing impaired (Mason & Kokkinakis, 2014; Reinhart et al., 2015).

## Acknowledgement

This work was supported by EU H2020-MSCA-RISE-2015 Grant No. 691229, VEGA 1/0355/20 and APVV-0452-12. EV was co-financed by Greece and the European Union (European Social Fund—ESF) through the Operational Programme «Human Resources Development, Education and Lifelong Learning» in the context of the project “Reinforcement of Postdoctoral Researchers—2nd Cycle” (MIS-5033021), implemented by the State Scholarships Foundation (IKY).

